# Genome-wide association study in New York *Phytophthora capsici* isolates reveals loci involved in mating type and mefenoxam sensitivity

**DOI:** 10.1101/2020.04.01.020826

**Authors:** Gregory Vogel, Michael A. Gore, Christine D. Smart

## Abstract

*Phytophthora capsici* is a soilborne oomycete plant pathogen that causes severe vegetable crop losses in New York (NY) State and worldwide. This pathogen is difficult to manage, in part due to its production of long-lasting sexual spores and its tendency to quickly evolve fungicide resistance. We single-nucleotide polymorphism (SNP) genotyped 252 *P. capsici* isolates, predominantly from NY, in order to conduct a genome-wide association study for mating type and mefenoxam insensitivity. The population structure and extent of chromosomal copy number variation in this collection of isolates were also characterized. Population structure analyses showed isolates largely clustered by the field site where they were collected, with values of F_ST_ between pairs of fields ranging from 0.10 to 0.31. Thirty-three isolates were putative aneuploids, demonstrating evidence for having up to four linkage groups present in more than two copies, and an additional two isolates appeared to be genome-wide triploids. Mating type was mapped to a region on scaffold 4, consistent with previous findings, and mefenoxam insensitivity was associated with several SNP markers at a novel locus on scaffold 62. We identified several candidate genes for mefenoxam sensitivity, including a homolog of yeast ribosome synthesis factor Rrp5, but failed to locate near the scaffold 62 locus any subunits of RNA Polymerase I, the enzyme that has been hypothesized to be the target site of phenylamide fungicides in oomycetes. This work expands our knowledge of the population biology of *P. capsici* and provides a foundation for functional validation of candidate genes associated with epidemiologically important phenotypes.

## Introduction

*Phytophthora capsici* is a soilborne oomycete plant pathogen that causes a disease known as Phytophthora blight on several economically important vegetable crops, including pepper, squash, and pumpkin. Phytophthora blight results in serious crop losses via both root and crown rot, which can lead to sudden irreversible wilting of the plant, and fruit rot, which manifests as rapidly expanding sunken lesions featuring a distinctive white sporulation (Granke et al. 2012). Vegetable growers have found *P. capsici* highly difficult to control for several reasons: it produces spores that can remain dormant in the soil for many years (Lamour and Hausbeck 2003, Carlson et al. 2017), rapidly reproduces asexually under suitable environmental conditions (Hausbeck and Lamour 2004), and tends to quickly evolve insensitivity to once effective fungicides (Parra and Ristaino 2001; Kousik and Keinath 2008).

Previous population genetic surveys of *P. capsici* isolates, both in Michigan (Lamour and Hausbeck 2001) and New York (NY; Dunn et al. 2010), have shown that different agricultural fields feature distinct, sexually recombining pathogen populations with limited gene flow between them. These results are consistent with the biology of *P. capsici*. Asexual sporangia and zoospores, both short-lived structures that cannot survive desiccation, are limited in their ability to spread quickly, as they are dispersed in water but not wind (Schlub 1983; Bowers et al. 1990; Granke et al. 2009). Sexual oospores, on the other hand, possess thick walls that allow them to survive harsh environmental conditions. Oospores can overwinter in temperate climates and remain viable in the soil for years (Bowers et al. 1990; Lamour and Hausbeck 2003; Babadoost and Pavon 2013; Carlson et al. 2017).

As a heterothallic species, *P. capsici* only reproduces sexually when isolates of opposite mating type are in contact (Erwin and Ribeiro 1998). The mating types of *Phytophthora* species, referred to as A1 and A2, do not signify the production of particular sex organs, as isolates of either mating type are hermaphroditic and thus capable of producing both antheridia and oogonia, the male and female reproductive organs, respectively (Ashby 1929; Judelson 1997). Rather, each mating type secretes a specific hormone (α_1_ or α_2_) that induces sexual reproduction in isolates of the opposite mating type (Ko 1978). Both mating type hormones have been isolated and characterized (Qi et al. 2005; Ojika et al. 2011) and mapping experiments using experimental crosses have identified a single locus controlling mating type in *P. capsici, P. infestans*, and *P. parasitica* (Fabritius and Judelson 1997; Lamour et al. 2012; Carlson et al. 2017). Researchers have hypothesized that mating type inheritance is analogous to sex determination in an XY system, where A1 isolates are homogametic (i.e., XX) and A2 isolates heterogametic (i.e., XY) at the mating type locus (Fabritius and Judelson 1997; Carlson et al. 2017). Nevertheless, the exact genes conferring mating type remain unknown.

Once a sexual population of *P. capsici* is established in a field, the eradication of the pathogen is highly unlikely and control strategies must be implemented to manage disease in future years. In addition to cultural practices designed to promote soil drainage and thereby deny a suitable environment for *P. capsici*, fungicides are one of the most effective means that growers rely on to control Phytophthora blight (Hausbeck and Lamour 2004). Phenylamide fungicides in particular, first the racemic metalaxyl and later its active enantiomer mefenoxam, have been used extensively since the 1970s due to their systemic activity in plants and high toxicity against many oomycete species via inhibition of ribosomal RNA (rRNA) synthesis (Davidse et al. 1983; Wollgiehn et al. 1984; Erwin and Ribeiro 1998). Shortly after these fungicides were first deployed, however, resistance emerged in populations of several important oomycete plant pathogens (Gisi and Sierotzki 2008). Isolates of *P. capsici* insensitive to mefenoxam were first reported in 1997 in New Jersey, North Carolina, and Michigan (Lamour and Hausbeck 2000; Parra and Ristaino 2001), and have since been found in many additional states, including NY (Keinath 2007; Wang et al. 2009, Dunn et al. 2010). As with mating type, the genes involved in mefenoxam insensitivity in *Phytophthora* species are largely unknown.

Once thought by many researchers to be haploid in their vegetative state, as are true fungi, cytological evidence in the 1960s and 70s established oomycetes as diploid with meiosis occurring in the gametangia immediately prior to fertilization of the oogonium (Sansome 1961; Sansome and Brasiere 1973). In recent years, data generated by next-generation sequencing technologies have been increasingly used not only for traditional population genetic analyses, but also for estimating ploidy levels of samples (Farrer et al. 2013; Zhu et al. 2016). As a result, researchers have identified polyploid *Phytophthora* isolates with three or more copies of each chromosome (Yoshida et al. 2013), as well as aneuploid individuals with a high degree of chromosomal copy number variation within a single genome (Barchenger et al. 2017; Shrestha et al. 2017). Nevertheless, there is still little known about the extent and distribution of chromosomal copy number aberrations within *P. capsici* field populations.

Since 2006, our lab has collected hundreds of isolates of *P. capsici* from vegetable-growing regions of NY State. We set out to leverage this collection in combination with isolates from an additional six states to improve our understanding of patterns of genomic variation in *P. capsici* and to discover loci associated with traits of epidemiological importance. To accomplish this, we genotyped over two hundred isolates from our culture collection with genotyping-by-sequencing (GBS) and assayed them for their mating type and mefenoxam sensitivity. The specific objectives of this study were to i) characterize the pathogen population structure in NY and compare findings with expectations based on previous inferences about pathogen dispersal and survival (Dunn et. al, 2010), ii) determine the extent of chromosomal copy number variation among isolates of *P. capsici* collected from the field, and iii) conduct a genome-wide association study (GWAS) to identify loci involved in the genetic control of mating type and mefenoxam sensitivity.

## Materials and Methods

### Isolate collection, phenotypic assays, and DNA extraction

Isolates described in this study were either recently collected in NY in 2017 or 2018 (*n*=157), obtained from stored cultures isolated from NY sites prior to 2017 (*n*=85), or received from out-of-state collaborators (*n*=10; Table 1). NY pathogen samples from 2017 and 2018 were isolated from plant parts of several different species with Phytophthora blight symptoms collected at farms and one supermarket (Table S1). Small pieces of surface-disinfected tissue were plated on PARPH medium (Jeffers and Martin 1986) except for samples with evident sporulation, in which case sporangia were directly transferred to the surface of PARPH Petri plates. Plates were then incubated at room temperature for one to two weeks before transferring a plug from the edge of the growing colonies to new PARPH plates. These plates were sealed with Parafilm (Bemis, Neenah, WI, USA) and stored at room temperature in the dark until used to transfer plugs for isolation of single-zoospore cultures.

**Table 1.** Number of isolates, unique genotypes, and counts of each mating type and mefenoxam sensitivity class per site. CD = Capital District, CNY = Central New York, LI = Long Island, WNY = Western New York, S = mefenoxam sensitive, IS = mefenoxam intermediately sensitive, R = mefenoxam resistant. Non-NY isolates were received from California (*n*=2), Florida (*n*=1), Michigan (*n*=1), New Mexico (*n*=1), Ohio (*n*=1), and South Carolina (*n*=4). Mefenoxam sensitivity counts for Ontario #1 isolates in 2013 are incomplete due to the loss of some cultures in storage prior to performing sensitivity assays.

| Field | Region | Year | Number<br>of Isolates | Mating Type |  | Mefenoxam |  |  | Number Unique<br>Genotypes |
| --- | --- | --- | --- | --- | --- | --- | --- | --- | --- |
|  |  |  |  | Counts |  | Sensitivity Counts |  |  |  |
|  |  |  |  | A1 | A2 | S | IS | R |  |
| Columbia #1 | CD | 2014 | 1 | 0 | 1 | 1 | 0 | 0 | 1 |
| Herkimer #1 | CD | 2007 | 1 | 1 | 0 | 0 | 1 | 0 | 1 |
| Rensselaer #1 | CD | 2007 | 2 | 1 | 1 | 1 | 0 | 1 | 2 |
| Schenectady<br>#1 | CD | 2007 | 2 | 2 | 0 | 1 | 1 | 0 | 2 |
| Schenectady<br>#2 | CD | 2007 | 1 | 1 | 0 | 0 | 0 | 1 | 1 |
| Cayuga #1 | CNY | 2017 | 24 | 9 | 15 | 20 | 3 | 1 | 10 |
| Monroe #1 | CNY | 2006 | 1 | 1 | 0 | 1 | 0 | 0 | 1 |
| Ontario #1 | CNY | 2013 | 70 | 34 | 36 | 44 | 1 | 0 | 17 |
| Ontario #1 | CNY | 2017 | 71 | 34 | 37 | 65 | 4 | 2 | 42 |
| Ontario #2 | CNY | 2018 | 9 | 0 | 9 | 8 | 1 | 0 | 2 |
| Ontario #3 | CNY | 2006 | 1 | 0 | 1 | 1 | 0 | 0 | 1 |
| Tioga #1 | CNY | 2018 | 1 | 0 | 1 | 1 | 0 | 0 | 1 |
| Tompkins #1 | CNY | 2018 | 1 | 1 | 0 | 1 | 0 | 0 | 1 |
| Suffolk #1 | LI | 2018 | 6 | 1 | 5 | 6 | 0 | 0 | 6 |
| Suffolk #2 | LI | 2018 | 4 | 1 | 3 | 4 | 0 | 0 | 2 |
| Suffolk #3 | LI | 2018 | 4 | 2 | 2 | 1 | 0 | 3 | 2 |
| Suffolk #4 | LI | 2018 | 3 | 3 | 0 | 2 | 0 | 1 | 3 |
| Suffolk #5 | LI | 2007 | 3 | 1 | 2 | 3 | 0 | 0 | 3 |
| Erie #1 | WNY | 2017 | 17 | 10 | 7 | 9 | 1 | 7 | 8 |
| Erie #2 | WNY | 2017 | 16 | 10 | 6 | 1 | 6 | 9 | 9 |
| Erie #3 | WNY | 2018 | 1 | 1 | 0 | 0 | 0 | 1 | 1 |
| Non NY | NA | NA | 10 | 5 | 5 | 8 | 1 | 1 | 10 |
| NY unknown | NA | NA | 3 | 2 | 1 | 3 | 0 | 0 | 3 |
| Total | NA | NA | 252 | 120 | 132 | 181 | 19 | 27 | 129 |

New York isolates collected prior to 2017, including nine described in previous publications (Dunn et al. 2010; Parada-Rojas and Quesada-Ocampo 2018; Table S1), were obtained from long-term storage tubes containing sterile water and 3-4 hemp seeds. Cultures of these isolates were started by plating a sample of the contents of the storage tubes onto PARPH plates. Cultures of non-NY isolates, including one previously described (12889; Foster and Hausbeck 2009), were received from collaborators and transferred to PARPH plates. All isolates in this study were verified as *P. capsici* by performing PCR with species-specific primers (Dunn et al., 2010) and confirming the presence of a band of the correct size via gel electrophoresis.

Single-zoospore isolates were obtained using a protocol similar to that described by Lamour and Hausbeck (2000). Isolates were induced to sporulate by plating on unclarified V8 agar and incubating under 14h daily fluorescent lighting for 7-14 days. Sporangia were then transferred with a sterile pipette tip to 1.5 mL microcentrifuge tubes containing 1 mL sterile water and incubated at room temperature for 45 min to promote zoospore release. Zoospore suspensions were serially diluted and 100 μl aliquots of 1:10, 1:100, and 1:1000 dilutions were spread-plated on water agar plates. After incubating at room temperature for approximately 16h, a stereo microscope was used to identify single germinating zoospores and transfer them to PARPH plates. Single-zoospore cultures were obtained for all but six isolates, which either featured poor colony re-growth (IMK328, FL29, 568OH) or inadvertently had DNA extracted prior to single-zoosporing (1070_3, 14_51, 2014_21).

Mating type and mefenoxam sensitivity assays were performed as described previously (Dunn et al. 2010). Briefly, for mating type determination, isolates were co-cultured on unclarified V8 agar plates with NY isolates of known mating type (A1: 0664-1 and A2: 06180-4) (Dunn et al. 2014). After incubating plates in the dark for 7-14 days, mating type was ascertained by confirming the presence or absence of oospores in the media using a stereo microscope. For a subset of isolates, consisting of 45 of the 2013 Ontario #1 isolates and 7 isolates from 2006-07 previously characterized by Dunn et al. (2010), mating type was determined at two distinct time points, both prior to entering and after removal from long term storage. When discordant, the mating type reported and used for analysis was the mating type determined at the time point tissue was collected for DNA extraction (corresponding to prior to long term storage for the 2013 Ontario #1 isolates and after removal from long term storage for the 2006-07 isolates).

Mefenoxam sensitivity was determined by transferring 10mm plugs from the edge of growing colonies on PARPH to unclarified V8 media amended with 5 μg/ml or 100 μg/ml mefenoxam (Ridomil Gold EC; Syngenta AG, Basel, Switzerland). After incubating plates in the dark for 3 days, colony diameters were measured and used to calculate percentage relative growth (RG) compared to control plates without mefenoxam added. These assays were repeated twice for each isolate and mean RG values are reported. In order to report a single summary statistic per isolate, isolates were classified as sensitive (<40% RG on 5 μg/ml mefenoxam), intermediately sensitive (>40% RG on 5 μg/ml mefenoxam but <40% RG on 100 μg /ml mefenoxam), or resistant (>40% RG on 100 μg/ml mefenoxam) as in Silvar et al. (2006).

Mycelia collection and DNA extraction were performed as previously described (Carlson et al. 2017). Plates containing 10% clarified V8 broth were inoculated with three plugs from the edge of an actively growing culture and incubated in the dark for 4-5 days. Approximately 100 mg of mycelia was then vacuum-filtered, collected into 2 ml microcentrifuge tubes containing two zinc-plated steel BBs (Daisy, Rogers, Arkansas, USA), and stored at −80°C prior to DNA extraction. DNA was extracted using the DNeasy Plant Mini Kit (Qiagen, Valencia, CA, USA) according to the manufacturer’s instructions, except a TissueLyser (Qiagen, Valencia, CA, USA) was used to disrupt mycelia and DNA was eluted into ultrapure water instead of AE buffer.

### Genotyping, SNP quality control, and clone correction

DNA samples were sent to one of two facilities to prepare and sequence 96-plex GBS libraries digested with *Ape*KI. Libraries for samples from the Ontario #1 field site from 2013 were prepared in 2015 at the Institute for Genomic Diversity at Cornell University and sequenced on an Illumina Hiseq2500 generating single-end 100 bp reads. Libraries for remaining samples were prepared in 2019 at the University of Wisconsin-Madison Biotechnology Center and sequenced on an Illumina NovaSeq6000 generating paired-end 150 bp reads. Replicate samples of isolate 0664-1 were included in order to assess technical variation in genotype calls between GBS plates.

Genotypes were called with the TASSEL GBSv2 pipeline (Glaubitz et al. 2014) using default parameters. To ensure that the same fragments were sequenced for all samples, only the forward reads were used for samples that were paired-end sequenced. Alignment of unique tags to the *P. capsici* reference genome (Lamour et al. 2012) and mitochondrial genome (provided by Martin, F., USDA-ARS) assemblies was performed with the Burrows-Wheeler algorithm bwa-aln (bwa version 0.7.17) with default parameters (Li and Durbin, 2009). Genotype calls were output in variant call format (VCF) and filtered initially for several criteria using VCFtools version 0.1.17 (Danecek et al. 2011). Indels and SNPs with >2 alleles were removed, as were individuals with greater than 60% missing data. SNPs that met the following criteria were then retained: 1) mean read depth >8 and <50; 2) >50% call rate; and 3) minor allele frequency (MAF) >0.01. Genotypes supported by fewer than 5 reads were set to missing. This SNP set was used for clonal group identification and determination of chromosomal copy numbers.

To identify clonal groups, pairwise identity-by-state (IBS), defined as the proportion of alleles shared at non-missing sites, was calculated among all isolates. We selected 95% as a threshold for declaring two isolates as clones, in order to account for an expected genotyping error rate of ^~^3% (Carlson et al. 2017), which was consistent with the error rate between technical replicates included among our samples. For all groups of isolates with pairwise IBS that surpassed this threshold, the isolate with the least missing data was retained to create a clone-corrected dataset, which was then further filtered to retain only SNPs with MAF >0.05 and call rate >80%. SNPs that were heterozygous in >80% of isolates were also removed, in order to remove likely artifacts of alignment errors. This clone-corrected SNP set was used for all population genetics and GWAS analyses.

All analyses, unless otherwise specified, were performed using custom R scripts (R Core Team 2019; https://github.com/gmv23/pcap-gwas). Deviations from a 1:1 mating type ratio among the clone-corrected isolate set for each site were tested using an exact binomial test with the R function *binom.test* at α=0.05. The percentage of variation in RG on mefenoxam-amended media attributed to differences between clonal lineages was calculated from the variance components of a mixed linear model predicting RG, with a random effect for clonal lineage and a fixed intercept term. Models were fit using the R package lme4 (Bates et al. 2015).

### Population structure

A principal component analysis (PCA) was conducted using the pcaMethods R package (Stacklies et al. 2007) on the unit-variance scaled and centered genotype matrix, and using the *nipals* method to account for missing data. The neighbor-joining tree was created in SplitsTree version 4.14.8 (Huson 1998), using the uncorrected “P” method to estimate the distance matrix, and plotted using the R package ape (Paradis et al. 2004). Values of pairwise F_ST_ between fields, as measured by Weir and Cockerham’s estimator of F_ST_ (Weir and Cockerham 1984), were calculated with the R package StAMPP (Pembleton et al. 2013).

### Chromosomal copy number determination

Because there is currently no chromosome-level genome sequence for *P. capsici*, we assigned a total of 97,145 SNPs to the 18 linkage groups of the genetic map provided in Lamour et al. (2012). We then used allele balances at heterozygous sites, defined as the number of reads of the major allele divided by the total number of reads, to identify putative linkage groups with more than 2 copies present in a given isolate, following methods similar to those of Farrer et al. (2013). To minimize variation in allele balances caused by low read depths, we only calculated allele balances at sites supported by 12 reads. For each isolate, SNP allele balances on linkage groups with 30 or more heterozygous SNPs were divided into bins of 47-53% (as expected for a chromosome with 2 copies) and 30-36% or 63-69% (as expected for a chromosome with 3 copies). We did not attempt to explicitly identify linkage groups that were present in four or more copies because of the ambiguity in expected allele balances (for example, an allele balance of 0.50 could indicate a copy number of either 2 or 4). Bootstrap analysis was performed 1000 times over sites and linkage groups were assigned copy number designations if they had over 95% bootstrap support for featuring predominantly disomic or trisomic allele balances.

Read depth variation across all sites was then used to confirm linkage group copy number designations for putative aneuploid isolates. For each of these isolates, the number of sites on each linkage group was balanced by downsampling to the number of sites on the linkage group featuring the fewest non-missing SNPs. We then performed an ANOVA to determine if site read depth varied across linkage groups and Tukey’s HSD to conduct pairwise comparisons across all linkage groups for a given isolate. Putative trisomic linkage groups only retained their trisomic designation if they featured an average read depth significantly different from all putative disomic linkage groups for that isolate at a family-wide confidence level of 0.05. Fisher’s exact test, using the R function *fisher.test*, was performed to test for significant enrichment for both trisomy across linkage groups and aneuploidy across clonal lineages.

### Linkage disequilibrium calculation and genome-wide association study

Pairwise linkage disequilibrium (LD), as measured by *r^2^* (Hill and Robertson 1968), was estimated between all SNPs within 500 kb of each other using the software program PopLDDecay version 3.40 (Zhang et al. 2019). Because RG on both 5 μg/ml and 100 μg/ml mefenoxam-amended media were not normally distributed, mefenoxam sensitivity was converted into a binary trait for the purpose of a better fitting model in GWAS (i.e. improved control of Type I error rate) using a threshold of >10% RG on 5 μg/ml mefenoxam to classify an isolate as resistant. We did not use the RG on 100 μg/ml mefenoxam phenotype as it was highly correlated (*r* = 0.95) with RG on 5 μg/ml mefenoxam.

A genome-wide association study for each phenotype was conducted using logistic regression models implemented in the R package GENESIS (Gogarten et al. 2019). All missing genotype calls were conservatively imputed with the mean value prior to conducting association tests. The Akaike information criterion (Sakamoto et al. 1986) was used to compare the fit of different generalized linear mixed models that included up to four PCs as covariates (to control for population structure) and a random polygenic isolate effect with a covariance structure defined by a kinship matrix (to control for unequal relatedness). The kinship matrix was estimated from the genotypic data using the *A.mat* function in the R package rrBLUP (Endelman 2011). In the mating type GWAS, we fit a generalized linear model for each SNP that only included a fixed intercept term, as PCs were not selected for inclusion in the model and the random isolate effect (i.e., kinship) was negligible. For mefenoxam sensitivity, SNPs were tested for association in a generalized mixed linear model that included an intercept term, the first three PCs retained as fixed effects, and a random isolate effect. SNPs with a Bonferonni-adjusted *P*-value <0.05 were considered significant. Manhattan and Q-Q plots were created using the R package qqman (Turner 2014).

### Data availability

Fastq files containing the raw, demultiplexed GBS reads were deposited in the National Center of Biotechnology Information Sequence Read Archive (accession number SRP101385; BioProject accession number PRJNA376558). Metadata on all isolates, including collection years and locations as well as mating type and mefenoxam sensitivity phenotypes, are provided in Table S1. Filtered VCF files for both the non-clone corrected and clone corrected isolate sets are available at CyVerse (link upcoming). Scripts used for analysis are available on Github (https://github.com/gmv23/pcap-gwas).

## Results

### Isolate collection

Between 2007 and 2018, 242 axenic *P. capsici* cultures were isolated from symptomatic plant samples collected at 22 field sites and a supermarket (Tompkins #1) in NY State (Figure 1). Geographic areas were represented disproportionately and were sampled in different years, with almost all of the Capital District isolates, for example, collected in 2007. The number of isolates collected from each site also varied considerably, with eleven sites each contributing 1 isolate and another site, Ontario #1, contributing 141 isolates (Table 1). Ontario #1 was the only site where samples were collected in two different years, with 70 isolates sampled in 2013 and 71 sampled in 2017.

**Figure 1.**
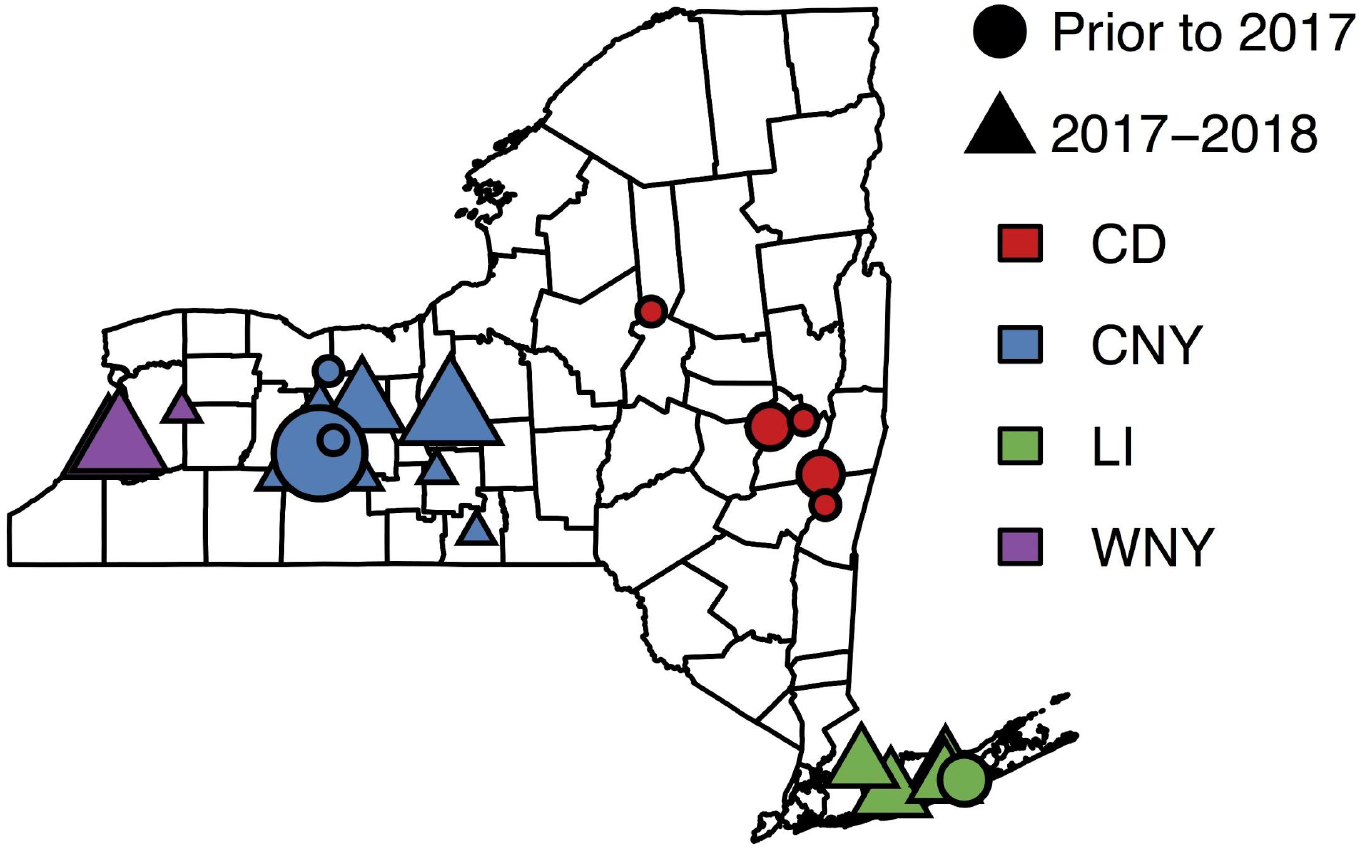
Map of NY sites where isolates were sampled. Points were randomly assigned a location within the counties where sites were located, with point size indicative of the number of isolates collected per site. One site in Ontario county was sampled twice, in both 2013 and 2017. CD = Capital District, CNY = Central New York, LI = Long Island, WNY = Western New York.

Isolates were found to belong to both mating types, with a total of 115 A1 and 127 A2 isolates collected (Table 1). A subset of the isolates were assayed for mating type twice, both shortly after collection and after re-culturing from long-term storage tubes, up to 13 years later. Of the 52 isolates assayed twice, 13 showed evidence of a mating type switch (Table S1). In all 13 of these cases, the mating type was A1 prior to storage and A2 upon re-culture.

Isolates that were intermediately sensitive or resistant to mefenoxam were identified in 12 of the 23 sites across the state (Table 1). Among the seven of these sites where multiple isolates were collected, Erie #2 featured the greatest proportion of intermediately sensitive or resistant isolates (94%).

### Genotyping and clonality

The 242 NY isolates and 10 isolates from other states (CA, FL, MI, NM, OH, and SC) were genotyped using GBS. From a total of 363,044 SNPs discovered, initial filtering criteria resulted in a dataset of 107,569 filtered SNPs called on 245 isolates, after removing 7 samples with high missing data. This SNP set, featuring a mean SNP read depth with a median of 17.4 calculated across sites and a median of 18.9 calculated across samples, was used to calculate pairwise IBS among isolates in order to identify isolates clonally derived from a common ancestor. Whereas the median IBS between all pairs of samples was 0.741, pairwise IBS between technical replicates of isolate 0664-1 (including samples sequenced by different facilities) ranged between 0.980 and 0.987. To account for genotyping error, we therefore used a 95% IBS threshold for declaring two isolates as clones. The 245 isolates were resolved into 129 unique genotypes (Table 1), 41 of which featured multiple isolates and are hereafter referred to as clonal lineages. A total of 157 isolates belonged to one of these clonal lineages, with a median clonal lineage size of three isolates, and the largest clonal lineage consisting of 17 isolates from Ontario #1 in 2013. The proportion of isolates that were genetically unique in each site varied, with Ontario #1 in 2013 (17 unique genotypes out of 70 isolates) and Ontario #2 in 2018 (2 out of 9) featuring the least genotypically diverse samples.

Each clonal lineage was private to a particular site, and in the case of Ontario #1, none of the clonal lineages identified in 2013 appeared again in 2017. Isolates belonging to the same clonal lineage were largely consistent in terms of mating type and mefenoxam sensitivity. Out of the 41 clonal lineages, only one – consisting of one A1 and two A2 isolates from Ontario #1 in 2013 – featured isolates of opposite mating type. Seven clonal lineages contained isolates with discordant mefenoxam sensitivity classifications, but these only featured a mixture of resistant and intermediately sensitive isolates (4 clonal lineages) or intermediately sensitive and sensitive isolates (3 clonal lineages). Among all isolates belonging to one of the 41 clonal lineages, 96.1% and 91.7% of the variation in RG on 5 μg/ml and 100 μg/ml mefenoxam, respectively, was attributed to differences between lineages.

A representative isolate was sampled from each clonal lineage to generate a clone-corrected data set consisting of 129 isolates. Using the clone-corrected isolate set, none of the site-years featured a ratio of mating types that differed significantly from 1:1, consistent with the neutral expectation for a sexually reproducing pathogen.

### Population structure

In order to visualize genetic relationships among the isolates, we conducted a PCA and created an unrooted NJ tree using a high-confidence set of 64,124 SNPs (Figure 2). Principal component 1, explaining 15.61% of the variance in the SNP genotype data, mainly separated the 59 isolates from Ontario #1 from the remaining isolates. Principal component 2 accounted for 4.5% of the variance explained and differentiated Western NY isolates from the 10 isolates sampled from Cayuga #1.

**Figure 2.**
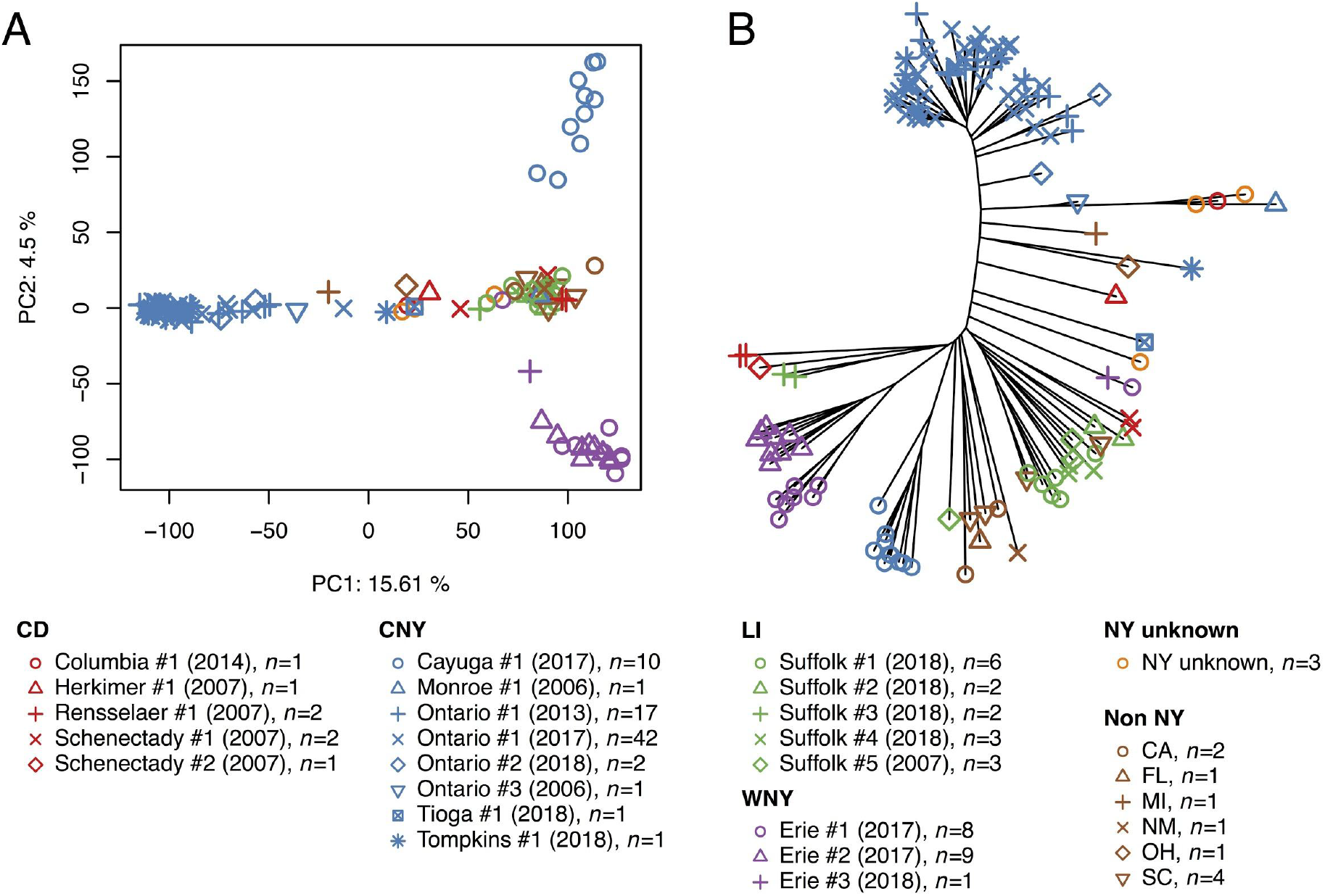
Population structure of the clone-corrected isolate set. A) Principal component analysis plot showing principal components 1 and 2 calculated from 64,630 SNPs scored on 129 genetically unique isolates. B) Unrooted neighbor-joining tree of the 129 isolates. In both plots, point color reflects geographic region and point shape reflects site within geographic region.

In the NJ tree, isolates from the same site were largely, but not exclusively, monophyletic. Exceptions included isolates from 3 of the 5 Long Island sites, as well as the Erie #1 and Suffolk #5 populations, each of which featured a single isolate appearing in a distinct clade. The outlier isolate from Erie #1, collected in 2017, appeared closely related to an isolate sampled from a distinct site in Erie county in 2018 (Erie #3). Sites from the same region grouped together in several cases, such as with Erie #1 and Erie #2 in Western NY and 4 of the 5 Long Island sites. However, isolates from different sites in the Capital District did not cluster together on the tree and Tompkins #1 was clearly separated from the remaining Central NY sites. Non-NY isolates were sorted into several different clades, with even isolates from the same state, such as the 4 isolates from South Carolina, clustering separately. In both the PCA and the NJ tree, no differentiation was observed between the 2013 and 2017 populations from Ontario #1.

To quantify differentiation between individual sites, we calculated pairwise F_ST_ for the 5 fields featuring >5 isolates after clone-correction (Table 2). Ontario #1 isolates from 2013 and 2017 were considered jointly as one population due to their genetic similarity (F_ST_=0.001). Values of F_ST_ ranged from a moderate differentiation of 0.10 (Erie #1 vs Erie #2) to a very strong differentiation of 0.31 (Cayuga #1 vs Ontario #1).

**Table 2.**
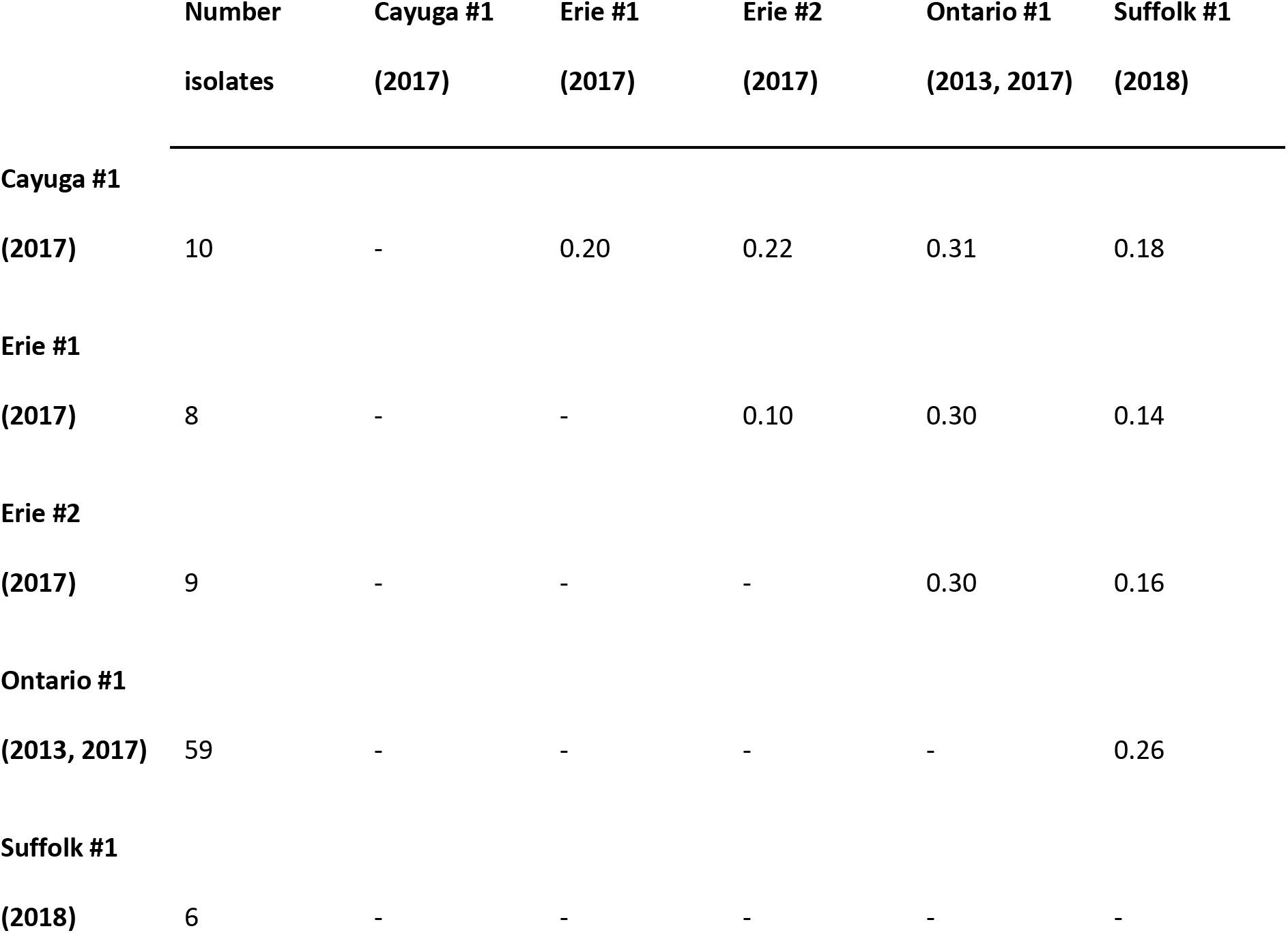
Estimates of Weir and Cockerham’s F_ST_ between sites featuring more than 5 isolates after clone-correction. Ontario #1 populations from 2013 and 2017 were collapsed into one population.

### Chromosomal copy number variation

Allele balances at heterozygous SNPs and intragenome variation in read depth were used to infer copy numbers for the 18 linkage groups in each of the 245 isolates with filtered genotype data (Figure 3; Table S2). For a majority of the isolates (*n* = 222), we were unable to assign a copy number to certain linkage groups because they either contained too few heterozygous markers or showed ambiguous signal in their allele balances or read depths (Figure S1). This noisy signal was likely caused, for the most part, by low average read depth, since individual mean read depth and the number of linkage groups assigned a copy number per isolate were highly correlated (*r*=0.79). Across isolates, the median number of linkage groups assigned a copy number was 13.

**Figure 3.**
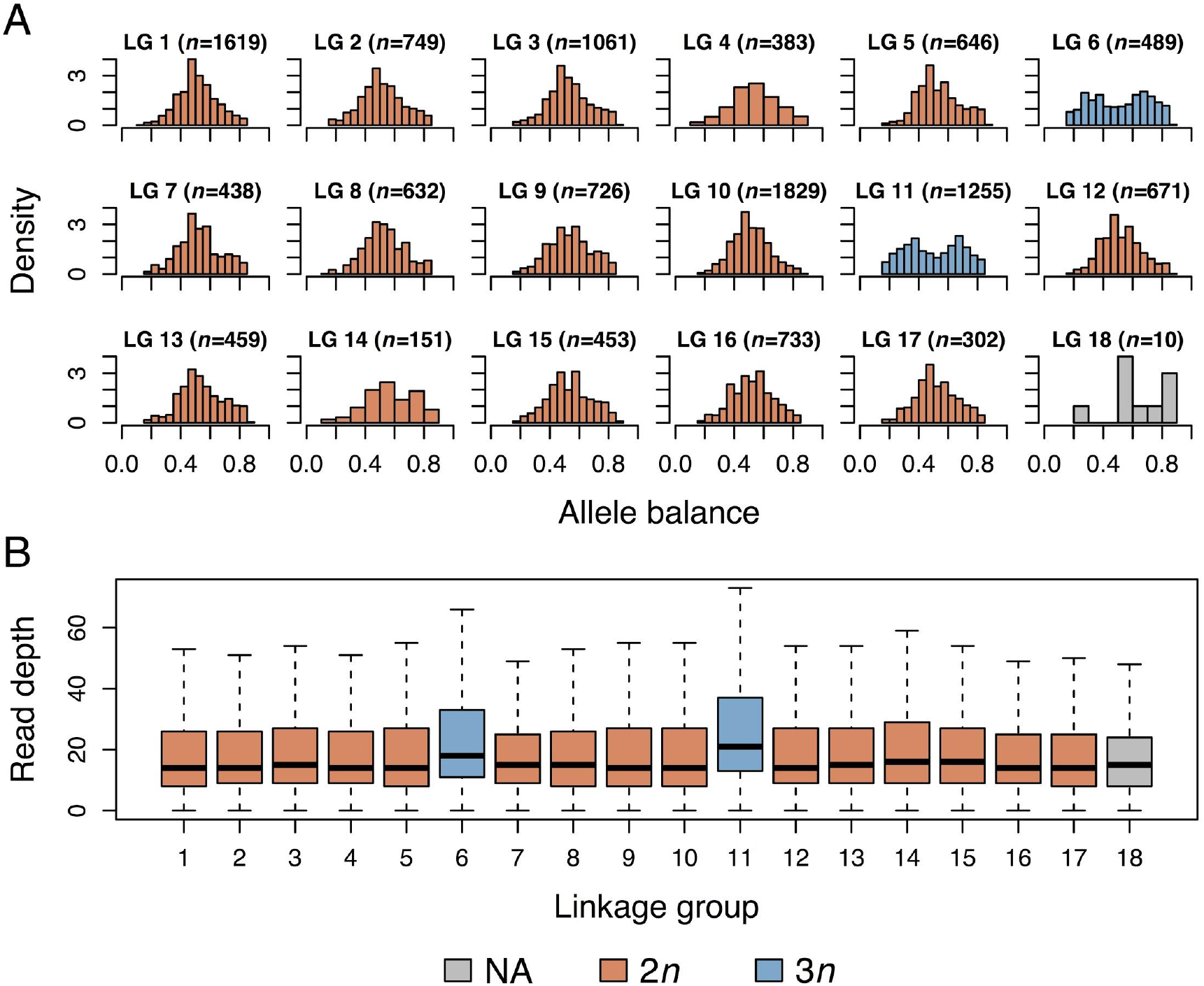
Example of ploidy level determination by linkage group in isolate 13EH38A. A) Distribution of allele balances within the 18 linkage groups of isolate 13EH38A. Number of heterozygous markers (*n*) is reported for each linkage group. B) Boxplot of SNP read depths per linkage group in isolate 13EH38A. The number of SNPs on each linkage group was downsampled to the number on the linkage group having the fewest SNPs.

In total, 31 isolates were aneuploid, having between 1 to 4 trisomic linkage groups. Linkage groups featuring allele balances suggestive of trisomy mostly featured higher read depths than putative diploid linkage groups, except for four isolates (FL29, 17EH33C, 1070_3, and A2_6_1) where trisomic linkage groups featured lower than average read depth, suggesting a possible tetraploid or higher base ploidy level. Two additional isolates, 17EH20A and 17EH21C, were triploid for all of their linkage groups whose copy number we could determine (13 and 16 linkage groups for 17EH20A and 17EH21C, respectively). Although these two isolates were the exclusive representatives of a particular clonal lineage, patterns of aneuploidy were not consistent within clonal lineages for the remainder of the isolates. In fact, among the 10 clonal lineages featuring at least one aneuploid isolate, all trisomic linkage groups were private to individual isolates. There was no significant enrichment for aneuploidy in any of these 10 clonal lineages compared to the rest (Fisher’s exact test *P*-value=0.35).

All linkage groups appeared trisomic in at least one isolate. Considering only linkage groups assigned a copy number designation, Fisher’s exact test showed a significant enrichment for trisomies in certain linkage groups compared to others (*P*=0.01), with linkage groups 6 and 17 displaying the highest rates of trisomy, appearing trisomic in 7 and 8% of isolates, compared to a linkage group average of 3% (Table 3).

**Table 3.** Distribution of copy number counts for each linkage group (LG) across 245 isolates. Reported are the number of isolates in which a particular LG showed strong statistical support for the presence of 2 copies (‘2*n*’) or three copies (‘3*n*’), as well as the number of isolates where that linkage group showed ambiguous signal (‘Unknown’) or featured too few heterozygous markers for analysis (‘NA’). The proportion of trisomic isolates per linkage group is also reported among isolates where that linkage group was assigned a copy number (‘Proportion 3*n*’).

| Linkage group | $2n$ | $3n$ | Unknown | NA | Proportion $3n$ |
| --- | --- | --- | --- | --- | --- |
| 1 | 214 | 2 | 27 | 2 | 0.01 |
| 2 | 174 | 3 | 65 | 3 | 0.02 |
| 3 | 179 | 3 | 60 | 3 | 0.02 |
| 4 | 156 | 3 | 82 | 4 | 0.02 |
| 5 | 185 | 3 | 54 | 3 | 0.02 |
| 6 | 138 | 11 | 91 | 5 | 0.07 |
| 7 | 101 | 2 | 138 | 4 | 0.02 |
| 8 | 191 | 4 | 47 | 3 | 0.02 |
| 9 | 168 | 2 | 72 | 3 | 0.01 |
| 10 | 208 | 3 | 31 | 3 | 0.01 |
| 11 | 186 | 4 | 52 | 3 | 0.02 |
| 12 | 161 | 4 | 77 | 3 | 0.02 |
| 13 | 152 | 3 | 87 | 3 | 0.02 |
| 14 | 130 | 2 | 108 | 5 | 0.02 |
| 15 | 130 | 4 | 107 | 4 | 0.03 |
| 16 | 178 | 4 | 60 | 3 | 0.02 |
| 17 | 131 | 11 | 99 | 4 | 0.08 |
| 18 | 57 | 3 | 110 | 75 | 0.05 |
| Sum | 2839 | 71 | 1367 | 133 | NA |

### Linkage disequilibrium decay and genome-wide association studies

We used phenotypes and genotypes from the clone-corrected isolate set and conducted GWAS to identify loci associated with mating type and mefenoxam sensitivity. First, we plotted LD between pairwise SNPs as a function of distance, finding that genome-wide LD decayed rapidly to *r^2^* < 0.10 by ^~^12 Kb, and plateaued at *r^2^* = 0.04 by ^~^400 Kb (Figure S2). We therefore focused on the 400 kb flanking regions of peak SNPs identified in GWAS for candidate gene searches.

Mating type showed no relationship with population structure, as it was not associated with any of the first four PCs (t-test *P*=0.60, 0.70, 0.90, 0.60 for PCs 1-4, respectively; Figure 4A). In addition, both mating types were represented approximately equally among the clone-corrected isolate set (n_A1_=66; n_A2_=63; Figure 4B). A logistic regression GWAS for mating type revealed one peak on scaffold 4 containing 70 SNPs with significant *P*-values (Figures 4C and 4D). The 70 significant SNPs spanned an approximately 426 Kb region (between SNPs S4_447285 and S4_873767; Figure S3) containing 139 annotated genes, with the peak SNP (S4_579765; *P*=4.3 × 10^−16^) located in an intron of a gene encoding a putative protein kinase (*fgenesh1_pg.PHYCAscaffold_4_#_142*). Because the molecular mechanism of mating type determination is currently unknown, we were unable to identify any strong candidate genes among the 290 annotated genes in the 400 Kb region flanking the peak SNP. Among the 70 significant mating type-associated SNPs, A1 isolates were predominantly homozygous, with a median frequency of 0.94 for homozygous genotype calls. A2 isolates, on the other hand, were predominantly heterozygous at these sites, with a median frequency of 0.70 for heterozygous genotype calls.

**Figure 4.**
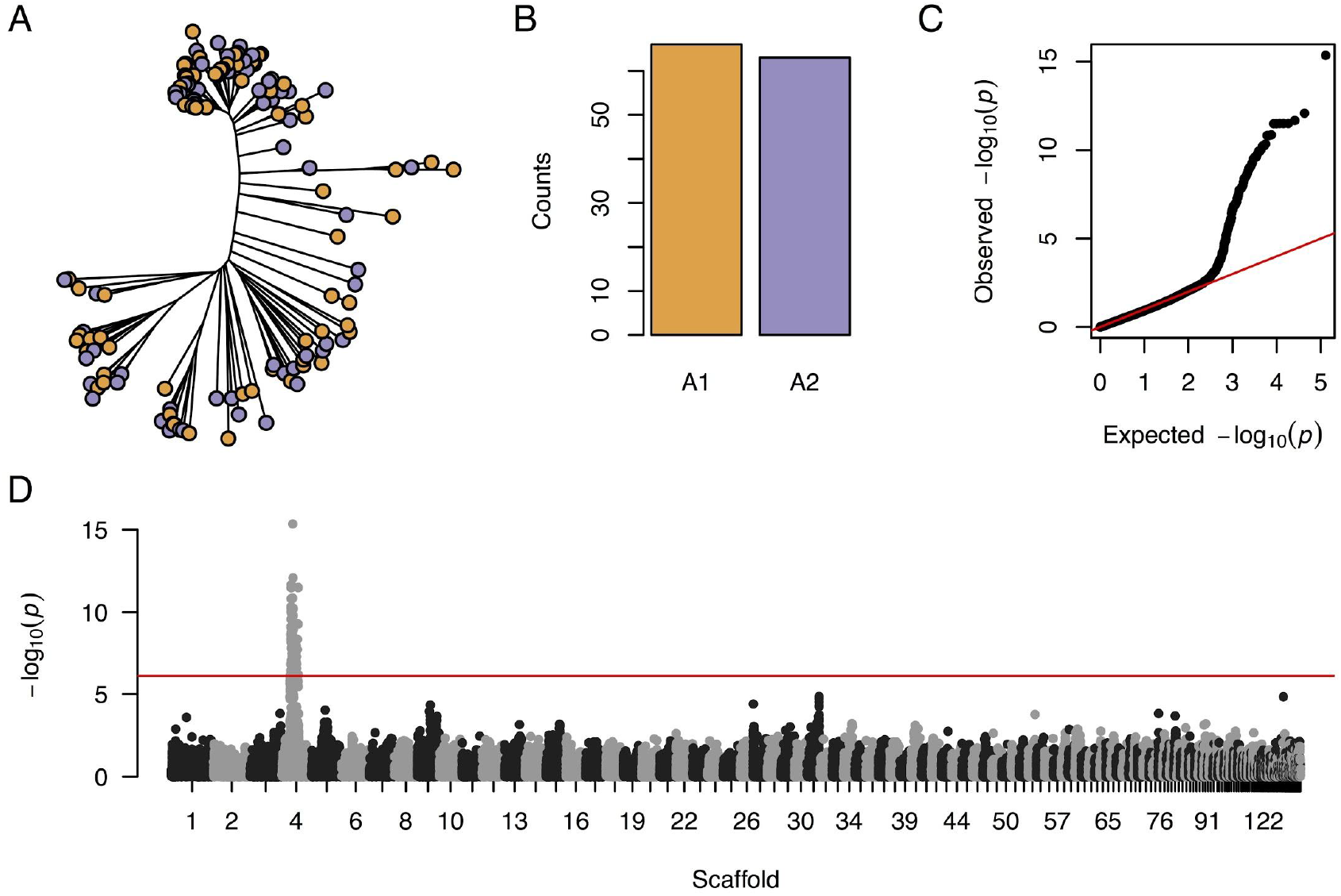
GWAS results for mating type. A) Neighbor-joining tree with point colors representing mating type where A1=orange and A2=purple. B) Counts of A1 and A2 isolates among the clone-corrected isolate set. C) Q-Q plot of *P*-values from logistic regression GWAS for mating type, using a population of 129 isolates scored for 64,630 SNPs. D) Manhattan plot showing SNP *P*-values across the genome from logistic regression GWAS for mating type.

We next investigated genotype discrepancies between isolates of the clonal lineage which featured one A1 (13EH26A) and two A2 (13EH05A and 13EH76A) isolates. While 13EH26A showed a genome-wide rate of discordant genotype calls between 13EH05A and 13EH76A of 4.3 and 4.7%, respectively, the discordant call rate was 36.7 and 36.5% in the 426 Kb area spanned by significant mating type-associated SNPs. Considering only the 70 significant SNPs, the discordant call rate was 100% and 97.7%. At 38 of the 42 significant SNPs without any missing genotype calls in these three isolates, 13EH05A and 13EH76A were both heterozygous and 13EH05A was homozygous. In addition to the mating type region on scaffold 4, 13EH26A also showed a higher rate of discordant genotype calls with 13EH05A and 13EH76A on scaffolds 34 and 40 (Figure S4), which immediately proceed the mating type region on the *P. capsici* genetic map (Lamour et. al. 2012). Collectively, these results suggest that a loss of heterozygosity event in a large chromosomal region containing the mating type locus induced a switch from A2 to A1 in isolate 13EH05A. Since the read depth of SNPs in 13EH05A in the mating type region (mean 23.1) did not differ from the genome-wide SNP read depth (mean 22.6), it appears that this loss of heterozygosity event was conferred in a copy-neutral manner.

In contrast to mating type, mefenoxam sensitivity was more strongly confounded with population structure, showing a significant association with PC 1 (*P*=5.9 × 10^−6^) and PC 3 (*P*=4.2 × 10^−3^). Intermediately sensitive and resistant isolates were especially prevalent in populations from WNY sites, and rare among the 59 isolates from Ontario #1 (Figure 5A). In addition, different phenotypic classes were disproportionately represented, with the majority of clone-corrected isolates classified as sensitive (Figure 5B). We used 10% RG on 5 μg/ml mefenoxam-amended media as a threshold to classify isolates as one of 34 resistant ‘cases’ or 91 sensitive ‘controls’ and conducted a logistic regression GWAS controlling for population structure and unequal relatedness. We identified one signal spanning approximately 59 Kb on scaffold 62 (between markers S62_139120 and S62_198531; Figure S4) that contained 6 significantly associated SNPs (Figures 5C and 5D). The peak SNP (S62_186715; *P*=1.1 × 10^−7^) was located in an exon of a gene (*gw1.62.46.1*) annotated as a dynein. The entirety of scaffold 62 was contained within the 400 Kb flanking the peak SNP and featured 100 annotated genes, including several with a plausible role in mefenoxam insensitivity, such as a homolog of the yeast ribosome synthesis factor Rrp5, located within 20 Kb of the peak SNP (Table S3).

**Figure 5.**
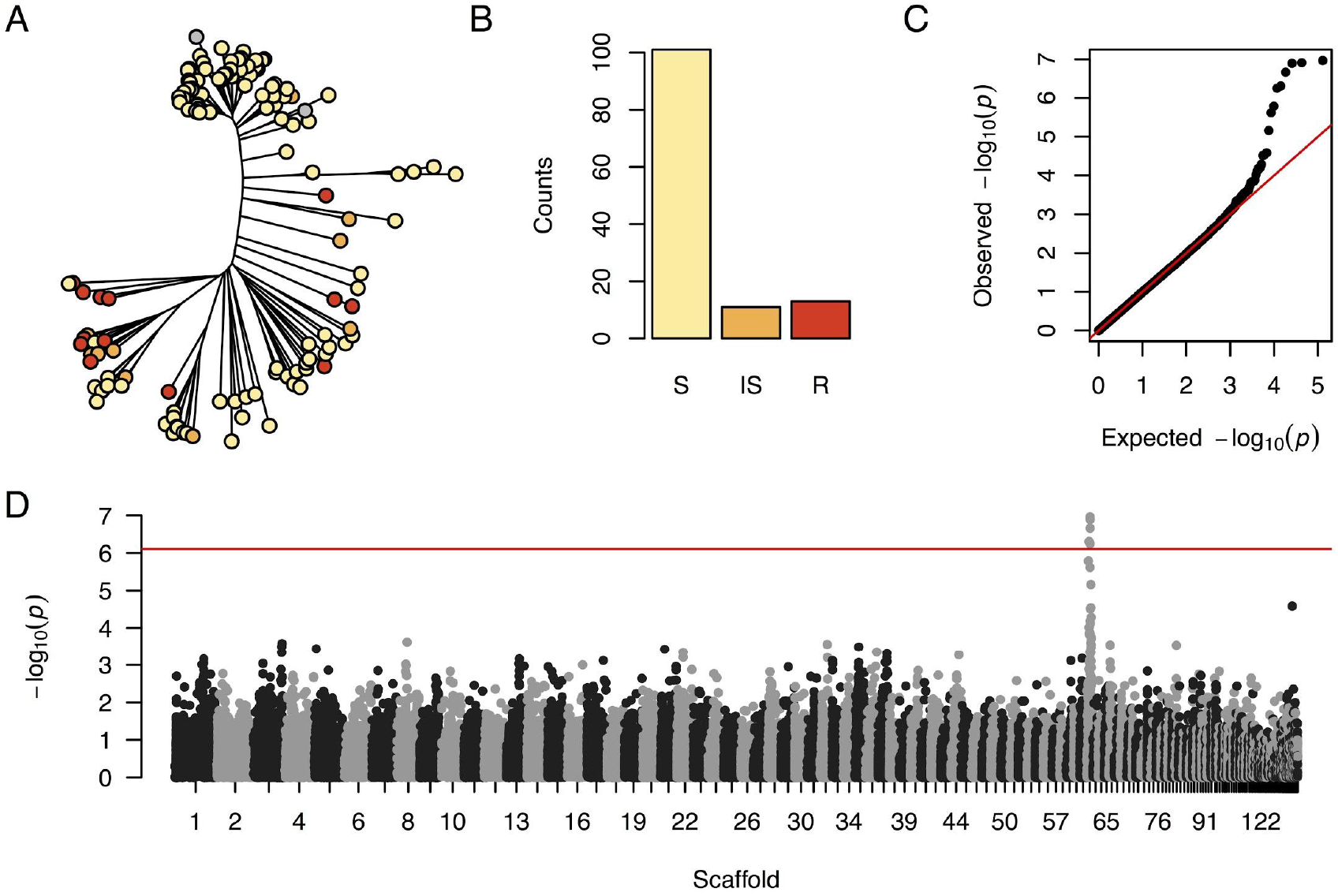
GWAS results for mefenoxam sensitivity. A) Neighbor-joining tree with point colors indicative of mefenoxam sensitivity. S = sensitive (yellow), IS = intermediately sensitive (orange), R = resistant (red). B) Counts of S, IS, and R isolates among the clone-corrected isolate set. C) Q-Q plot of *P*-values from logistic regression GWAS controlling for population structure and unequal relatedness. GWAS was conducted using a population of 125 isolates scored for 64,630 SNPs. D) Manhattan plot showing SNP *P*-values across the genome from logistic regression GWAS for mefenoxam sensitivity.

The allele associated with insensitivity at the peak SNP, S62_186715, showed an approximately additive effect on RG on both 5 μg/ml and 100 μg/ml mefenoxam (Figure 6). The insensitivity allele, with a frequency of 0.28 among all isolates, was found in isolates from 11 NY sites and in one SC isolate. Eight isolates, representing five different field sites in four NY regions, were homozygous for the insensitive allele. Several isolates were outliers in terms of their sensitivity response as predicted by the genotype of SNP S62_186715, including one isolate from Erie #1 that was homozygous for the sensitive allele yet showed >95% RG on 5 μg/ml mefenoxam and >75% RG on 100 μg/ml mefenoxam, as well as an isolate from Suffolk #5 that was homozygous for the insensitive allele yet did not grow on either 5 μg/ml or 100 μg/ml mefenoxam.

**Figure 6.**
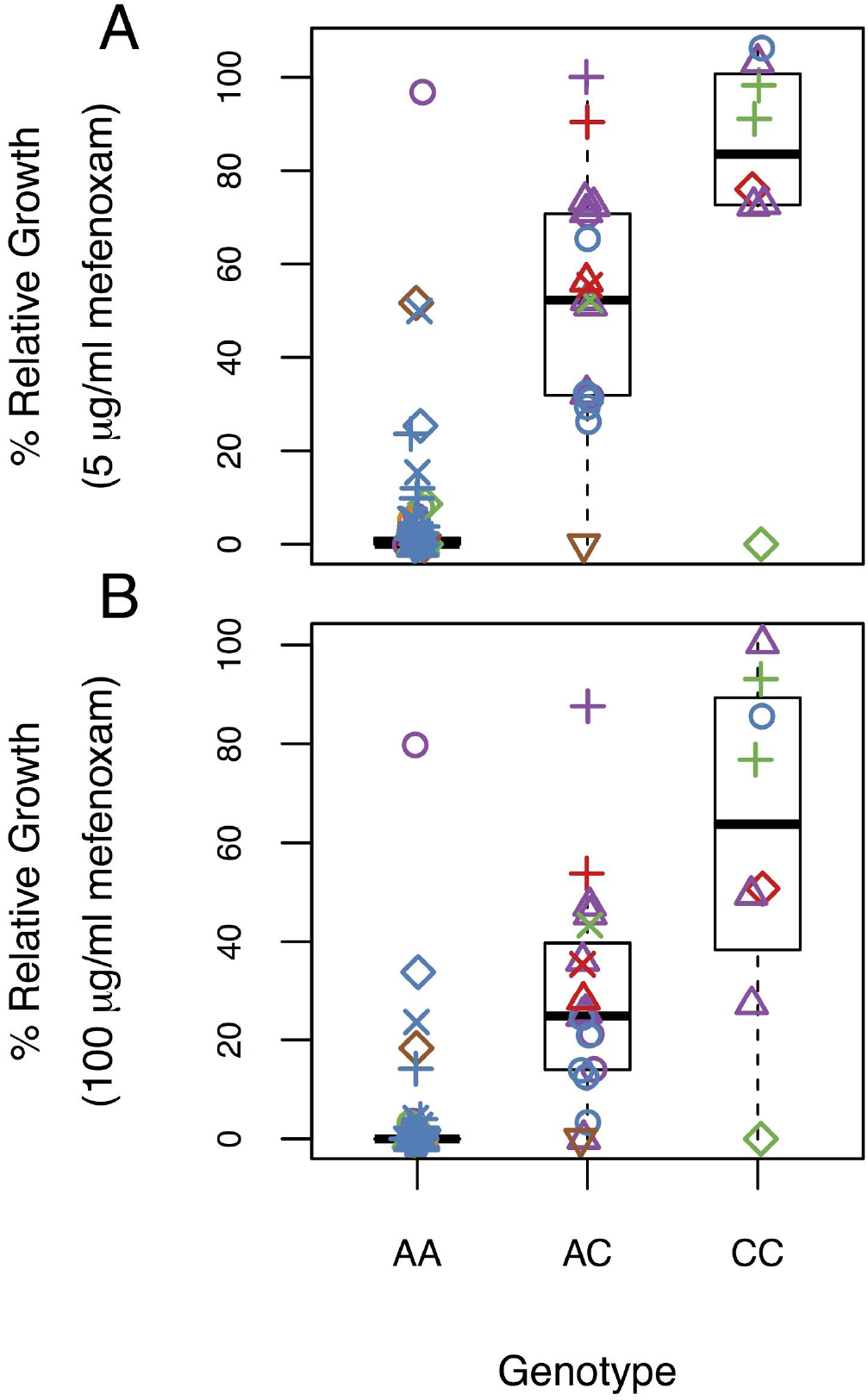
Boxplot showing the effect of the genotype at the peak SNP for mefenoxam insensitivity (S62_186715) on RG on amended media. Point colors and shapes are as in Figure 2. A) Effect of S62_186715 genotype on RG on 5 μg/ml mefenoxam. B) Effect of S62_186715 genotype on RG on 100 μg/ml mefenoxam.

## Discussion

The 252 *P. capsici* cultures characterized in this study were isolated from symptomatic plant samples obtained through several means – whether collected during field visits or received from extension agents for purposes of disease diagnostics. We genotyped this collection of isolates at 107,569 SNP loci, corresponding to one SNP on average every ^~^600 bp. We were able to leverage this dataset to investigate not only issues of regional importance, namely those related to the pathogen’s population structure patterns in NY, but also questions of global significance, having to do with the genetic control of mating type and mefenoxam sensitivity, both epidemiologically important phenotypes, and the extent to which field isolates vary in terms of their chromosomal copy numbers.

### Population structure and clonality

Visualizing relationships among the clone-corrected isolates with PCA and a NJ-tree (Figure 2) showed, for the most part, pathogen populations from different farms clustering separately. Similarly, estimates of pairwise F_ST_ between the five fields that featured a sufficient number of samples after clone-correction showed moderate to strong genetic differentiation between sites. Even isolates from farms in close geographic proximity, such as Erie #1 and Erie #2, separated into distinct, although related, clades in the NJ-tree and featured moderate levels of genetic differentiation between them. These results agree with previous hypotheses that limited gene flow occurs between *P. capsici* populations on different farms (Lamour and Hausbeck 2001, Dunn et. al. 2010).

Nevertheless, we found some examples that deviated from this overall trend. Isolates from 3 of the 5 Long Island farms did not separate into distinct clades in the NJ-tree, suggesting higher gene flow between the sampled fields in this region. We also identified two isolates, one each from Erie #1 and Suffolk #5, that did not cluster with other isolates from their respective sites. It is possible that these outlier isolates represent independent introductions of inoculum to their fields compared to the rest of the isolates from those sites.

As expected based on our current understanding of *P. capsici* epidemiology (Lamour and Hausbeck 2001, Lamour and Hausbeck 2003, Dunn et al. 2010), we found evidence for a mixed mode of reproduction in pathogen populations in NY fields. Over half of the isolates that we surveyed belonged to a clonal lineage consisting of multiple sampled isolates, reflecting the influence of asexual reproduction within single seasons and fields. Two fields, Ontario #1 (2013) and Ontario #2 (2018), featured a particularly small number of genetically unique isolates compared to the total number of isolates sampled from those fields (Table 1). Interestingly, within a few months prior to sample collection, both of these fields experienced flooding events followed by their first outbreak of Phytophthora blight. One possible explanation for their low observed genotypic diversity is that the inoculum, likely introduced to these fields via floodwater, consisted of a small number of founding pathogen genotypes. Alternatively, the environmental conditions in the fields after flooding, highly conducive to asexual reproduction, could have amplified the ability of certain clonal lineages to rapidly spread and predominate in the field, either due to random chance or because of a fitness advantage of those genotypes.

Isolates from Ontario #1 were sampled again in 2017, at the end of a season in which the field was planted for the first time since 2013 to a Phytophthora-susceptible crop and subsequently suffered another disease outbreak. The 2017 isolates, which were genetically undifferentiated from the 2013 isolates (F_ST_=0.001; Figure 2), were more genotypically diverse, with 59% of isolates representing unique genotypes compared to 24% in 2013. Given that the primary inoculum for the 2017 outbreak was most likely oospores formed during the 2013 outbreak, these results show how a potentially small inoculum founder event followed by sexual reproduction can lead to the development of a persistent bank of genotypically diverse oospores in the soil. This phenomenon has been previously demonstrated in both natural populations (Lamour and Hausbeck 2003) and experimentally inoculated research fields (Dunn et al. 2014, Carlson et al. 2017).

It is unknown when *P. capsici* inoculum first arrived in vegetable-growing areas of NY, nor where it originated from, or how many times it was independently introduced. Our population structure analyses did not show a clear relationship between NY isolates and those from any particular other state (Figure 2), although our non-NY sample size (*n*=10) was too modest to directly address this question. However, other studies have also shown only moderate differentiation between *P. capsici* populations from most U.S. states (Quesada Ocampo et al. 2011, Parada-Rojas and Quesada-Ocampo 2018), suggesting perhaps a recent common origin of inoculum for most states or frequent movement of inoculum between distant locations in the U.S. Our data, interpreted in conjunction with other studies (Lamour and Hausbeck 2000, Lamour and Hausbeck 2001, Lamour and Hausbeck 2002, Dunn et al. 2010, Jones et al. 2014) and reports from farmers, are consistent with a model of disease spread where local dispersal of inoculum, perhaps largely via run-off from fields and flooding events, leads to long-lasting sexual populations on newly infected fields that become genetically differentiated from neighboring populations due to a founder effect and subsequent genetic drift. Longer-distance movement of inoculum, mediated by transport of infested soil or infected plant-material, may also occur, as evidenced by our observation of several field populations with little relatedness to other populations in the same region (e.g. Cayuga #1).

Two observations from our collection of isolates – that A1 and A2 isolates were represented proportionally in all sites, and that clonal lineages were entirely unique to individual sites and years – reflect the epidemiological importance of sexual reproduction in allowing inoculum to overwinter. While obligate sexual reproduction appears to be the case in NY and other U.S. regions (Lamour and Hausbeck 2000, 2003), locations in South America and East Asia, including regions of China with temperate climates that should prohibit overwintering of asexual inoculum, have reported asexual lineages of *P. capsici* that persist over many years and over large geographic regions (Hurtado-Gonzalez et al. 2008, Hu et al. 2013). It is unknown what environmental and/or genetic factors account for the stark differences in the pathogen population structure between the U.S. and these other countries.

### Chromosomal copy number variation

By analyzing patterns of variation in read counts across the genome, we identified some isolates with clear evidence of aneuploidy (Figure 3), as well as others with much more difficult-to-interpret signal (Figure S1). We therefore decided to implement conservative thresholds for declaring a linkage group trisomic, relying on significance testing (whether parametric or bootstrap-based) with two independent sources of information, allele balances at heterozygous sites and total read depth at all SNPs. Low average read depth, causing noisy allele balance histograms, was likely responsible for many of the linkage groups with ambiguous support for an allele balance mode at either 0.5 or 0.33/0.66, as evidenced by the high correlation between individual read depth and the percentage of linkage groups assigned a copy number assignations in each isolate. However, it is also possible that linkage groups in some isolates showed noisy allele balance histograms for one of several biological reasons, whether due to possessing an even higher copy number than 3, having intra-chromosomal copy number variation, or because DNA was extracted from a heterogenous population of nuclei varying in copy number for that linkage group. We therefore may have underestimated the true extent of aneuploidy in our collection of isolates, and future efforts with higher sequencing depth are necessary to expand upon our research. It is also worth mentioning that some of the linkage groups we declared trisomic may actually have been present in greater than 3 copies, as higher-copy number linkage groups could feature allele balance patterns consistent with our definition of trisomy (for example, the tail of an allele balance peak at 0.75, as would be expected for a triplex tetraploid genotype, could cause a greater number of SNPs to have an allele balance closer to 0.66 than 0.50). Finally, our analyses do not rule out heterokaryosis (the presence of multiple distinct nuclei within a single cell), which was recently shown in the oomycete plant pathogen *Bremia lactucae* to result in similar allele balance distributions to those that we found (Fletcher et al. 2019). However, whereas *B. lactucae* produces multinucleate sporangia that germinate directly, we have observed *Phytophthora capsici* sporangia to almost exclusively germinate indirectly from mononucleate zoospores (data not shown), making heterokaryosis unlikely.

We found 33 isolates, representing 13% of the isolates we genotyped, that were either aneuploids or genome-wide polyploids. Other studies have reported both widespread polyploidy (Daggett et al. 1995) and aneuploidy (Barchenger et al. 2017, Shrestha et al. 2017) in isolates of *P. capsici* and other *Phytophthora* species. However, the number of isolates we genotyped and our detailed analyses allowed us to make several novel findings. First, we found a slight, but significant, enrichment for trisomy in certain linkage groups compared to others. It is unclear if this is due to chromosome size, since the physical size of the *P. capsici* chromosomes is unknown, or has to do with the presence of adaptive genes on certain linkage groups that confer a fitness advantage when present in higher copy number. Second, analyzing patterns of aneuploidy and polyploidy within clonal lineages showed evidence for a meiotic origin of polyploidy, as the only two genome-wide polyploids we found were the exclusive members of the same clonal lineage, and a mitotic origin of aneuploidy, since trisomic linkage groups were not shared in any cases between isolates of the same clonal lineage. Spontaneous chromosomal loss and duplication in *Phytophthora* species has previously been identified during vegetative growth and asexual reproduction (Kasuga et al. 2016, Hu et al. 2020), supporting aneuploidy arising mitotically. In our study, differences in aneuploidy within isolates of the same clonal lineage, which presumably originated from a single oospore germinating the same year as sample collection, suggest that chromosomal copy number variation arises rapidly. However, we cannot separate chromosomal duplications that occurred in the field from those that occurred in culture. Future efforts sequencing pathogen DNA directly from field samples would be necessary to understand the rate at which aneuploidy arises and under what conditions.

Aneuploidy is known to be either lethal or causative of severe development defects in many higher organisms (Siegel and Amon 2012). Nevertheless, under selective conditions, experimental evolution studies have shown that aneuploid yeast strains often have higher fitness than euploids (Sunshine et al. 2015). Indeed, in the Sudden Oak Death pathogen *Phytophthora ramorum*, aneuploid isolates with phenotypic alterations are consistently selected for after passage through certain hosts (Kasuga et al. 2016). Because of the small number of isolates in our collection having aneuploidy for any given linkage group, we were not able to robustly associate chromosomal copy number changes with either of the traits we phenotyped. However, we hypothesize that spontaneous chromosomal loss and duplication leads to phenotypic variation, even within clonal lineages of *P. capsici*, that are subject to selection. It is unclear, however, if selection for certain aneuploidies persist beyond a single year, given that *P. capsici* undergoes obligate sexual reproduction in order to overwinter in the United States and the ability of aneuploid isolates to undergo meiosis and form viable gametes is unknown.

### Genetic basis of mating type and mefenoxam sensitivity

To our knowledge, our study represents the first GWAS conducted in *Phytophthora capsici*, and one of two ever conducted in a plant-pathogenic oomycete (Ayala-Usma et al. 2019). LD decayed rapidly, to *r^2^* <0.10 by approximately 12 Kb, almost identical to the rate reported in a population of *P. infestans* (Ayala-Usma et al. 2019), and comparable to rates reported in three plant-pathogenic fungi: *Parastagonospora nodoroum, Zymoseptoria tritici*, and *Pyrenophora teres* (Gao et al. 2016, Hartmann et al. 2017, Anke et al. 2019). Background levels of LD (*r^2^*=0.04), however, were only reached at approximately 400 Kb, indicating the need for a fairly large candidate gene search space in GWAS.

We expected mating type to be easy to map via GWAS, since it is known to have a simple genetic architecture, represented a binary phenotype with both classes evenly distributed in our population, and was unassociated with population structure (Figure 4). Indeed, we found strong statistical support for a single mating type locus on scaffold 4 (Figure 4, Figure S3), the same region identified in previous studies (Lamour et al. 2012, Carlson et al. 2017). We were not able to identify strong candidate genes in this region due to our lack of knowledge of the molecular mechanism of mating type determination. It is unknown if the mating type locus comprises a single gene or several that are tightly linked, and the causal genes could presumably encode any of a wide variety of proteins, such as enzymes in the metabolic pathways for α_1_ or α_2_, receptors involved in the recognition of α hormones, or transcription factors responsible for regulating mating type-specific expression levels.

Previous evidence from experimental crosses supports a genetic model where mating type inheritance behaves as in an XY sex determination system, where A2 isolates are the heterogametic (i.e., XY) type and A1 isolates are the homogametic type (i.e., XX; Fabritius and Judelson 1997, Carlson et al. 2017). Our results are consistent with this hypothesis, as A1 isolates were predominantly homozygous and A2 isolates predominantly heterozygous at the 70 SNPs significantly associated with mating type. Other researchers have observed the A2 mating type to be unstable (Hu et al. 2013, 2020), with evidence suggesting that mitotic loss of heterozygosity in the mating type region causes A2 isolates to switch mating type (Lamour et al. 2012). While we did not identify any A2 isolates that became self-fertile, a phenomenon that was recently reported (Hu et al. 2020), we did observe 13 A2 isolates that converted to A1 at an undetermined point over a period of several years. In addition, we identified mating type discrepancies within a single clonal lineage, consisting of one A1 and two A2 isolates. While the three isolates were genetically identical across most of the genome, the A1 isolate featured a high rate of discordant genotype calls compared to the A2 isolates at the mating type locus and linked genomic regions (Figure S4), consistent with a loss of heterozygosity event in this isolate resulting in the conversion of the A2-determining haplotype. Because read depths were not halved in the mating type region compared to the rest of the genome, we hypothesize that this loss of heterozygosity occured in a copy-neutral manner such as gene conversion, as opposed to via a deletion event resulting in hemizygosity. Given what appears to be the high frequency of unidirectional switching from A2 to A1, it may be possible in a future experiment to fine-map the mating type locus by tracking loss of heterozygosity breakpoints in a collection of A2 isolates that have switched mating type.

Unlike mating type, mefenoxam sensitivity had never been mapped in *P. capsici* prior to thi*s* study. It also represented a more challenging phenotype to map, as it was confounded with population structure and exhibited an imbalance in the number of resistant versus sensitive isolates (Figure 5). Nevertheless, by fitting a generalized mixed model correcting for population structure and unequal relatedness, we detected significantly associated SNPs in a region on scaffold 62 (Figure 5, Figure S3). The allele associated with insensitivity at the peak SNP demonstrated an approximately additive effect (Figure 6), consistent with segregation patterns in lab crosses that support an incompletely dominant gene conferring metalaxyl or mefenoxam insensitivity in *P. capsici, P. infestans*, and *P. sojae* (Shattock 1988, Bhat et al. 1993, Gisi and Cohen 1996, Lamour and Hausbeck 2000). It appears likely, however, that additional loci may play a role in decreased sensitivity in some isolates, as several individuals in our collection of isolates were homozygous for the sensitive allele at the peak SNP on scaffold 62, yet showed intermediate or high mefenoxam insensitivity. Similarly, in *P. infestans*, there is evidence of multiple loci involved in insensitivity to metalaxyl (Judelson and Roberts 1999, Fabritius et al. 2007).

Researchers have long assumed that the target site of phenylamide fungicides is RNA Polymerase I (Griffith et al. 1992), since the mode of action of these chemicals in oomycetes involves inhibition of rRNA synthesis (Davidse et al. 1983, Wollgiehn et al. 1984). In *P infestans*, however, attempts to associate mutations in subunits of RNA Polymerase I with metalaxyl or mefenoxam insensitivity have been inconclusive (Randall et al. 2014, Matson et al. 2015). In our case, we did not identify any genes in the region of the GWAS signal on scaffold 62 encoding RNA Polymerase I subunits. We did, however, identify several other plausible candidate genes (Table S3). One gene, located within 15 kb of the peak SNP, is a homolog of yeast protein Rrp5, which is required for processing of pre-rRNA transcripts into the cleaved molecules that form the ribosome (Venema and Tollervey 1996). If mefenoxam does interfere with the function of Rrp5, the buildup of precursor rRNA molecules in the nucleolus could cause a depletion of total rRNA as these intermediate molecules are degraded by exonucleases or as feedback mechanisms result in decreased rRNA transcription. Alternatively, the causal gene at this locus could be one of the transporters located nearby, or perhaps even RNA polymerase III subunit Rpc4, also located within 15 Kb of the peak SNP. Further investigation is required to determine which of these genes, if any, feature mutations that confer mefenoxam insensitivity in the isolates we assayed.

### Conclusions

We genotyped a collection of NY *P. capsici* isolates at over 100,000 SNP markers and assayed them for their mating type and mefenoxam sensitivity. Results of population structure analyses were consistent with previous reports, showing limited gene flow between different fields and highlighting the importance of sexual reproduction in allowing inoculum to overwinter. Thirteen percent of the isolates we genotyped showed some degree of chromosomal copy number variation within their genomes, with linkage groups 6 and 17 featuring a particularly high rate of aneuploidy. Genome-wide association studies confirmed previous results mapping the mating type locus to a region on scaffold 4, and identified a novel locus associated with mefenoxam sensitivity on scaffold 62. These results provide a foundation for functional validation of candidate genes as well as molecular marker development for prediction of mating type and mefenoxam insensitivity. Furthermore, the panel of isolates we assembled and the dense marker data we generated represent a genetic resource that can be used for mapping of other important traits in *P. capsici*, such as sensitivity to additional chemicals or virulence on economically important host plants.

## Acknowledgements

This research was supported by the New York State Department of Agriculture and Markets (grant number C00237GG), the USDA National Institute of Food and Agriculture Specialty Crop Research Initiative (2015-51181-24285), and HM Clause, who provided support for Summer Research Scholars at Cornell AgriTech. We would like to thank Ali Cala, Santiago Tíjaro Bulla, Kimberly D’Arcangelo, Holly Lange, Charlotte Mineo, and Carolina Puentes Silva, all of whom provided valuable technical assistance in this project. We also thank Maryn Carlson, who collected a portion of the data and provided feedback and edits on an earlier version of this manuscript, as well as Di Wu and Dan Ilut, who offered bioinformatic assistance. In addition, we are grateful to Traven Bentley, Kishor Bhattarai, Elizabeth Buck, Mary Hausbeck, Brian Hill, Shaker Kousik, Margaret McGrath, Abby Seaman, and Darcy Telenko for providing cultures of isolates characterized in this study.

---

Table S1. Metadata on all isolates characterized in this study, including their source, mating type, mefenoxam sensitivity classification, and presence in the clone-corrected dataset.

Table S2. Copy number estimates for every linkage group in every isolate. 2*n* = statistical support for presence of 2 copies, 3*n* = statistical support for presence of 3 copies, ? = ambiguous copy number designation, NA = insufficient heterozygous markers on the linkage group for analysis.

Table S3. Curated list of candidate genes, along with their KOG and IPR annotations (Lamour et al. 2012), within the range of linkage disequilibrium (LD) decay of peak SNP associated with mefenoxam insensitivity. Distance to peak SNP (bp) was calculated as the physical distance of the peak SNP position from the open reading frame (ORF) start position.

Figure S1. Example of ploidy level determination by linkage group in isolate 12889MIA. A) Distribution of allele balances within the 18 linkage groups of isolate 12889MIA. Number of heterozygous markers (*n*) is reported for each linkage group. B) Boxplot of SNP read depths per linkage group in isolate 12889MIA. The number of SNPs on each linkage group was downsampled to the number on the linkage group having the fewest SNPs.

Figure S2. Pairwise linkage disequilibrium (*r*^2^) between SNPs as a function of physical distance (kb).

Figure S3. SNP *P*-values for association with mating type, represented by vertical lines, and linkage disequilibrium (*r*^2^) with the peak SNP, represented by triangles. This plot shows the region bounded by 400 Kb on either side of the peak SNP. The orange triangle represents the peak SNP associated with mating type.

Figure S4. Percentage discordant genotypes in non-overlapping 50 Kb bins in pairwise comparisons between isolates 13EH05A, 13EH26A, and 13EH76A. The three isolates are members of the same clonal lineage, yet 13EH05A and 13EH76A are of the A2 mating type and 13EH26A is A1. Only the first fifty scaffolds are shown for simplicity, with scaffold 45 not included because it did not have any SNPs.

Figure S5. SNP *P*-values for mefenoxam insensitivity, represented by vertical lines, and linkage disequilibrium (*r*^2^) with the peak SNP, represented by triangles, on scaffold 62. The orange triangle represents the peak SNP associated with mefenoxam insensitivity.

