## Supplementary material for "Genome-wide association study in New York *Phytophthora capsici* isolates reveals loci involved in mating type and mefenoxam sensitivity": Table S3 and Figures S1-S5

Table S1 and Table S2 (see excel documents).

Table S3.

| Gene ID | Protein ID | ORF start position (bp) | ORF stop position (bp) | Distance to peak SNP (bp) | KOG definition | IPR definition |
|---|---|---|---|---|---|---|
| fgenesh1_pg.PHYCAscaffold_62_#_1 | 20330 | 22463 | 26716 | -164252 | Pleiotropic drug resistance proteins (PDR1-15), ABC superfamily | ABC transporter-like;ABC transporter-like;CDR ABC transporter;AAA+ ATPase, core;ABC transporter-like |
| fgenesh1_pg.PHYCAscaffold_62_#_8 | 20337 | 110345 | 108642 | -76370 | NA | Biopterin transport-related protein BT1;MFS general substrate transporter |
| estExt2_fgenesh1_pm.C_PHYCAscaffold_620005 | 530348 | 114899 | 119014 | -71816 | Pleiotropic drug resistance proteins (PDR1-15), ABC superfamily | ABC transporter-like;ABC transporter-like;ABC-2 type transporter;AAA+ ATPase, core;ABC transporter-like |
| e_gw1.62.86.1 | 126238 | 142989 | 141670 | -43726 | Predicted transporter | Major facilitator superfamily MFS-1;MFS general substrate transporter |

| | | | | | | |
|---|---|---|---|---|---|---|
| e_gw1.62.160.1 | 126230 | 144559 | 143202 | -42156 | Predicted transporter | Major facilitator superfamily MFS-1;Major facilitator superfamily;MFS general substrate transporter |
| estExt2_fgenesh1_pg.C_PHYCAscaffold_620022 | 536757 | 175009 | 174164 | -11706 | DNA-directed RNA polymerase III subunit | RNA polymerase III Rpc4 |
| estExt2_Genewise1Plus.C_PHYCAscaffold_620123 | 554474 | 205528 | 204392 | 18813 | rRNA processing protein Rrp5 | RNA-processing protein, HAT helix |
| e_gw1.62.85.1 | 126220 | 221210 | 219951 | 34495 | Predicted transporter | Major facilitator superfamily MFS-1;Major facilitator superfamily;MFS general substrate transporter |
| fgenesh2_kg.PHYCAscaffold_62_#_28_#_Contig1951.1 | 510557 | 250135 | 249076 | 63420 | Uncharacterized membrane protein, predicted efflux pump | Multi antimicrobial extrusion protein MatE |

Figure S1

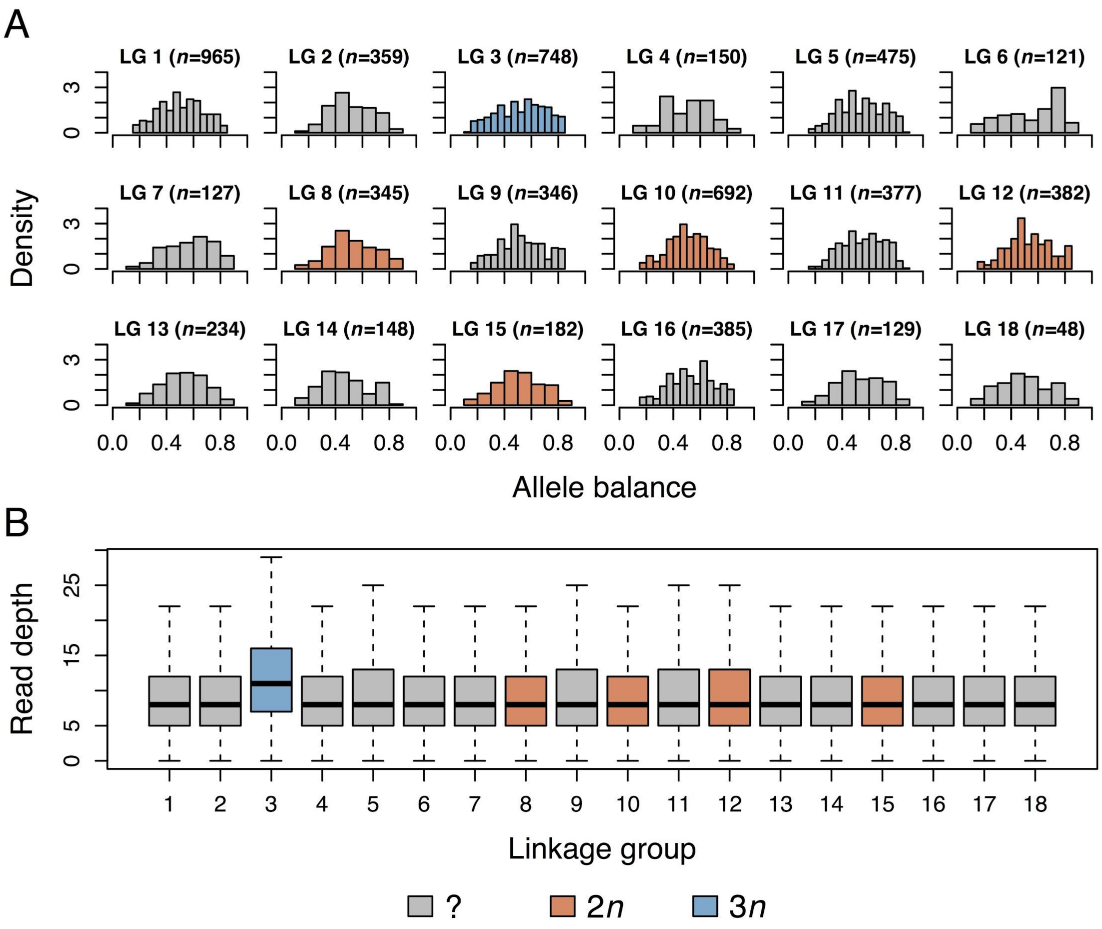
A
LG 1 (n=965)
LG 2 (n=359)
LG 3 (n=748)
LG 4 (n=150)
LG 5 (n=475)
LG 6 (n=121)
LG 7 (n=127)
LG 8 (n=345)
LG 9 (n=346)
LG 10 (n=692)
LG 11 (n=377)
LG 12 (n=382)
LG 13 (n=234)
LG 14 (n=148)
LG 15 (n=182)
LG 16 (n=385)
LG 17 (n=129)
LG 18 (n=48)
Density
Allele balance
B
Read depth
Linkage group
?
2n
3n

Figure S2

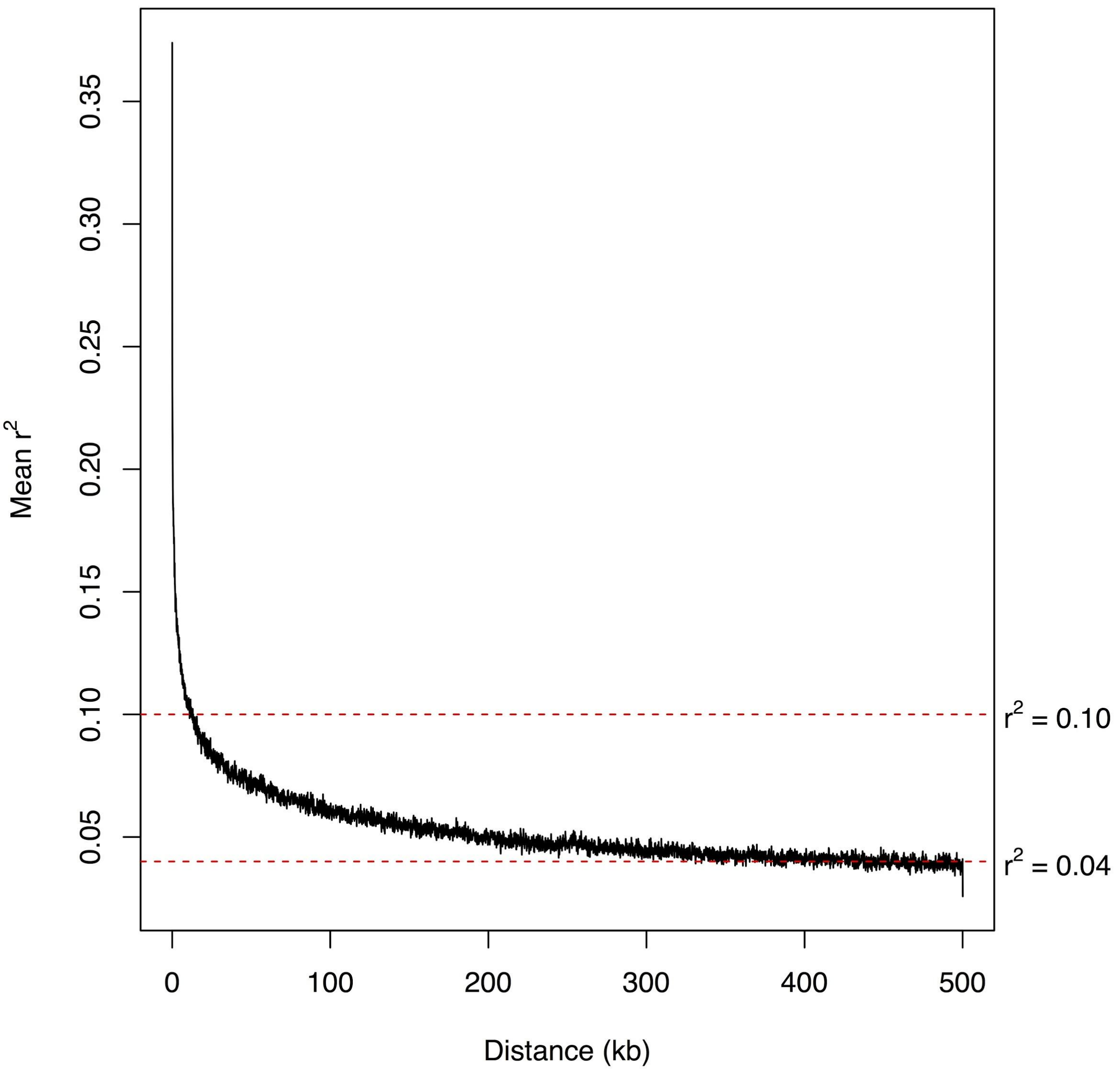

Mean $r^2$
0.35
0.30
0.25
0.20
0.15
0.10
0.05
$r^2 = 0.10$
$r^2 = 0.04$
0
100
200
300
400
500
Distance (kb)

Figure S3

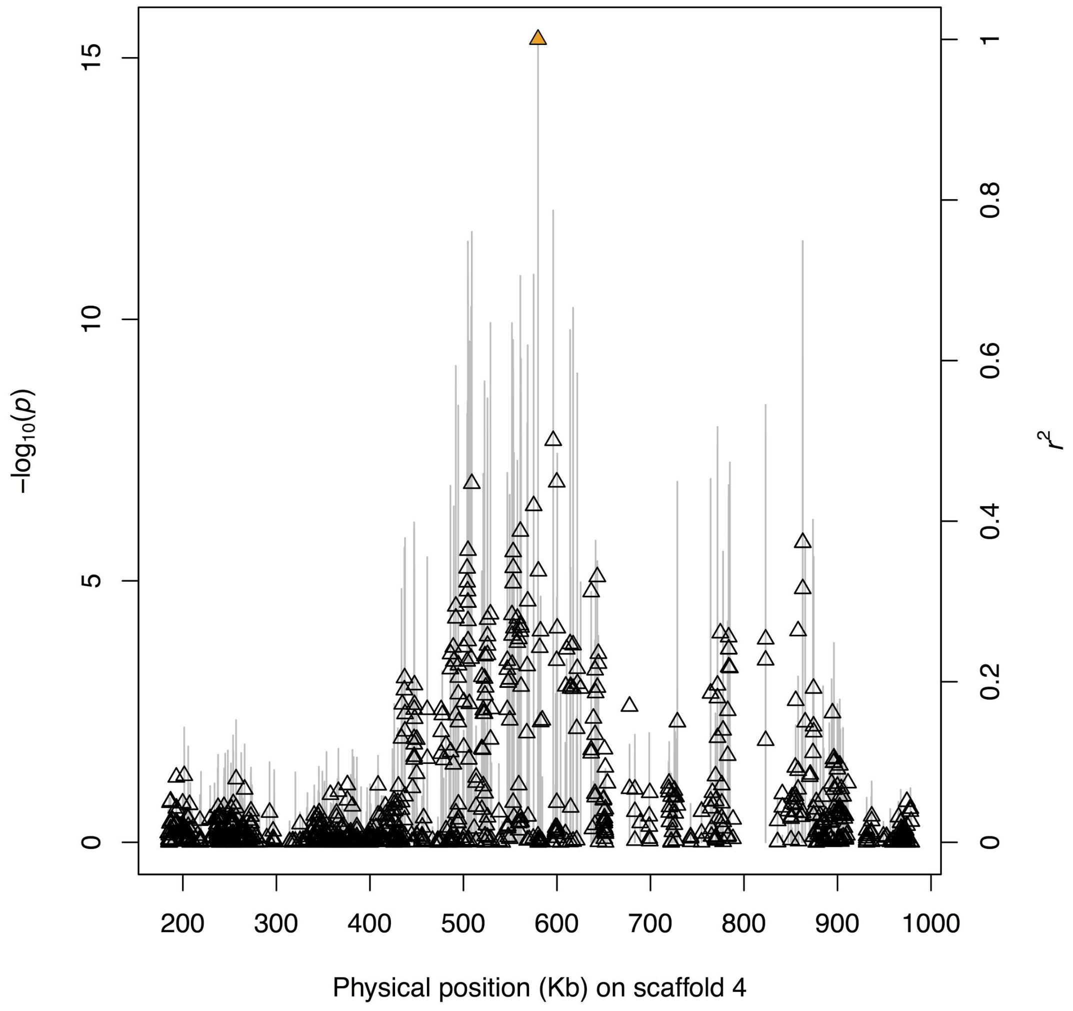

15
10
5
0
−log10(p)
1
0.8
0.6
0.4
0.2
0
r2
200
300
400
500
600
700
800
900
1000
Physical position (Kb) on scaffold 4

Figure S4

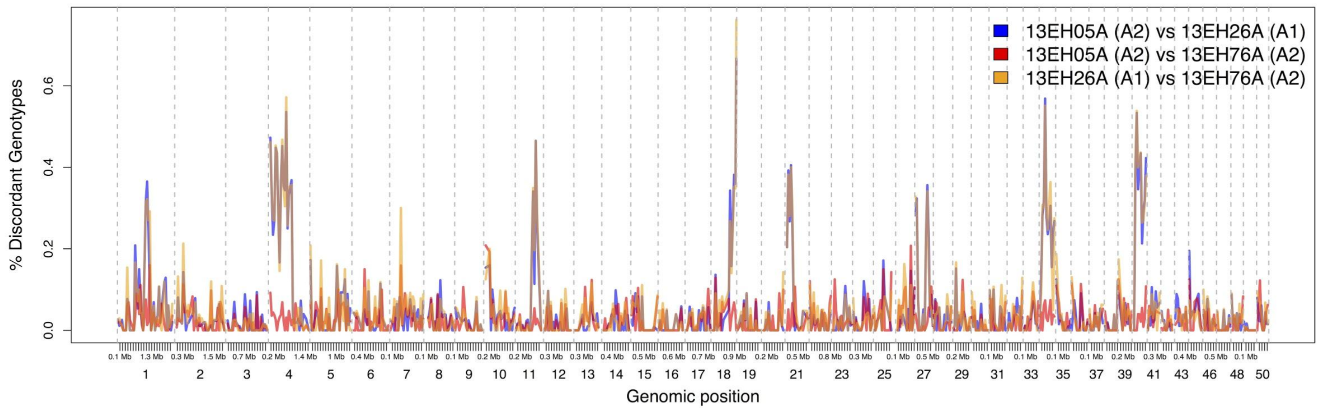


Figure S5

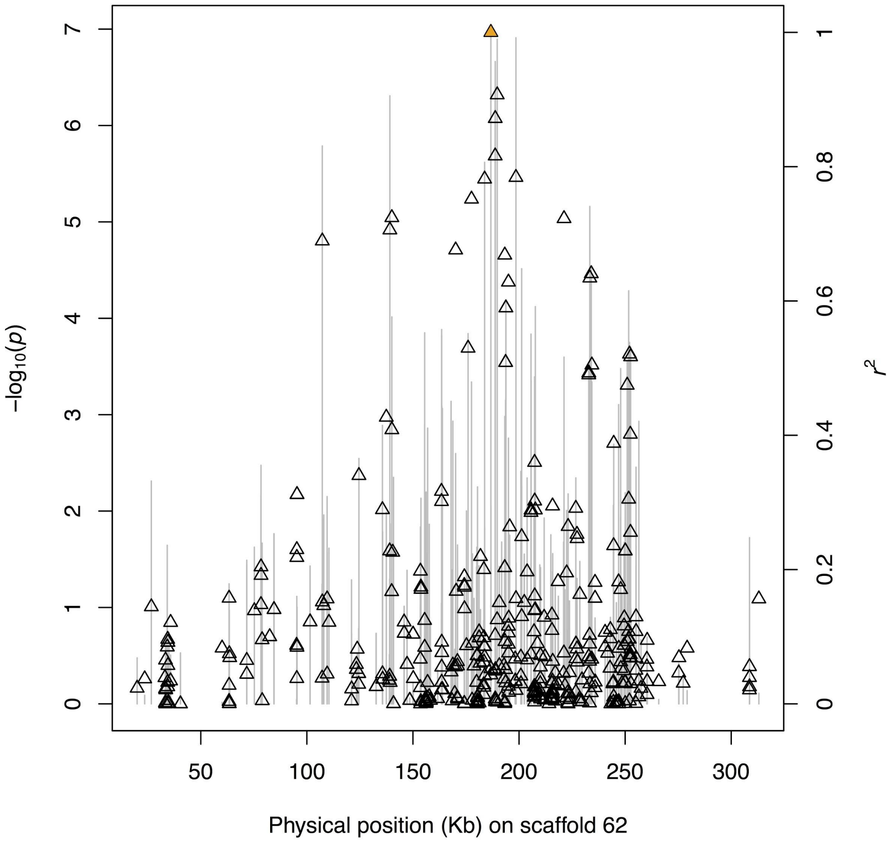

-log10(p)
r2
Physical position (Kb) on scaffold 62
0
1
2
3
4
5
6
7
50
100
150
200
250
300
0
0.2
0.4
0.6
0.8
1
